# ER shaping proteins guide spindle elongation and division during rapid cleavage mitoses

**DOI:** 10.64898/2026.07.15.738729

**Authors:** Katherine R. Rollins, Austin R. Clark, Prakriti Kandel, Schuyler B. van Engelenburg, J. Todd Blankenship

## Abstract

The ER is a complex network of membranes that inhabits much of the cytoplasm of cells – however, this network undergoes a massive condensation and rapid remodeling during cell division. In *Drosophila* cleavage divisions, this results in a tight association of the ER with centrosomes and mitotic spindle poles. Previous work has shown that this relationship between the ER and centrosomes must be finely tuned to enable successful spindle elongation, and that overaccumulation of the ER in these stages can result in failed centrosome maturation. During interphase, the ER exists in tubular and sheet-like arrangements, with a variety of “shaping” proteins enforcing these topologies. Here, we examine the contributions of these ER shaping proteins to the rapid changes that occur during cleavage mitoses in the *Drosophila* embryo. A screen of ER shaping proteins revealed that disruption of Reep-family proteins leads to mitotic failures at characteristic cleavage stages. Compromising *ReepA*, the *Drosophila* ortholog of the Reep1-4 subfamily, had a lesser impact on early embryonic mitoses. However, *ReepB* (the ortholog of the Reep5-6 subfamily) disruption, significantly affects ER mitotic coat morphologies, resulting in a ‘frilled ER’ phenotype and a reduction of ER adherence to the spindle space accompanied by division failures. Overexpressing ReepA does not rescue ReepB mitotic or ER morphology defects and instead introduces local condensates of abnormal ER structures. These data suggest that dedicated Reep proteins guide ER mitotic properties at specific early developmental stages. Using a cell-based *in vitro* analysis of *Drosophila* Reeps, we identify differential “tubulating” properties of ReepA and ReepB. Together these data suggest that the minutes-scale ER remodeling required for early mitoses is governed by shaping proteins, and that ReepB family members are especially important in some of the most rapid cleavage divisions that occur in early embryo.

## Introduction

Early development of multicellular organisms often necessitates rapid rounds of mitotic division to establish a population of cells that is large enough to support the development of complex tissue types and morphologies. These divisions, called the cleavage divisions, are specialized such that they can occur in rapid succession and entire mitotic cycles may take place in as little as a few minutes (Schejter and Wieschaus, 1993; Schindler-Johnson and Petridou, 2024). Accompanying these division processes is the remodeling and partitioning of a variety of subcellular compartments that will support cell function (reviewed in Carlton et al., 2020). One organelle that displays remarkable dynamics during mitosis is the endoplasmic reticulum (ER). In many different model systems, the ER has been shown to undergo mitotic transformations that include condensation of ER structures toward division poles as well as alterations in the amount of tubular and sheeted structures (Terasaki et. al., 1986; Bobbinec et al., 2003; Lu et. al., 2009; Puhka et. al., 2012; Wang et. al., 2013; Bergman et al., 2015). This polar accumulation of ER has largely been viewed as a mechanism that ensures the proper division of ER into the resultant daughter cells (Bobinnec et. al., 2003; Kors and Schlaitz, 2024). However, recent work in the *Drosophila* embryo has shown that misregulation of ER remodeling can affect mitotic activities including the key processes of centrosome maturation and spindle force generation (Araújo et al., 2022; Rollins and Blankenship, 2023). Whether structural changes in the ER during mitosis modulate centrosome and spindle function is less clear. ER shaping proteins, such as Reticulons, have been shown to accumulate at spindle poles, suggesting highly structured ER membranes may be present in these regions (Karabasheva and Smyth 2019), and mammalian Reep3/4 proteins have been shown to shape mitotic ER tubules (Kumar et al., 2019). Shaping proteins have also been implicated in organizing an ER-free space for chromosomal alignment during mitosis (Schlaitz et al., 2013). However, how these functions impact spindle function is unclear. Here, we use the *Drosophila melanogaster* syncytial embryo as a model system for understanding the structural rearrangements of the ER that enable high-fidelity mitoses on a rapid timescale characteristic of cleavage cycles.

Cleavage divisions are characterized by fast, and often reductive, cell cycles that swiftly increase the number of cells post-fertilization. In early *Drosophila* embryos, these cleavage cycles begin as nuclear divisions within the yolk of the syncytium. As divisions continue, these nuclei migrate axially (during cell cycles 1-4), then toward the periphery of the embryo (during cell cycles 5-9), to establish a syncytial blastoderm, in which nuclei organize just below the embryonic cortex (Blake-Hedges and Megraw, 2019). The final four syncytial division cycles (cell cycle 10 through 13) are considered the cortical divisions and are differentiated from the first nine cycles by the formation of transient plasma membrane furrows. These transient furrows serve to separate mitotic figures and anchor the spindle apparatus (Foe and Alberts 1983; Cao et. al., 2010; Holly et al., 2015; Mavor et al., 2016; Schmidt and Grosshans, 2018; Tillery et. al., 2018; Xie and Blankenship, 2018; Tam and Harris, 2024). The *Drosophila* embryo at these stages is highly enriched in lipids and possesses an endoplasmic reticulum that is gradually depleted with each round of new furrow formation (Rollins and Blankenship, 2023).

The ER is generally categorized as having two major shapes: membranous ER sheets and ER membrane tubules (Zhang and Hu, 2016). Typically, the perinuclear ER, which is contiguous with the nuclear envelope, is recognized as the Rough ER (RER), as it is studded with ribosomes necessary for protein synthesis. The RER is commonly associated with ER sheets (Shibata et al., 2006) and proteins that are resident to this perinuclear domain such as Ribosome binding protein 1 (RRBP1) have been implicated in ER sheet stabilization (Shibata et al., 2010). During mitosis, the ER undergoes a sheet-to-tubule transformation (Puhka et. al., 2012) suggesting a potential importance and/or dominance of tubular shaped ER in mitotic cells and embryos. The peripheral ER, which is located closer to the plasma membrane, is considered to possess increased tubular morphologies and forms a complex network (Shibata et al., 2006). Current understanding of tubule promoting proteins began with the characterization of Reticulons (Rtns) and DP1/Yop1 proteins (Voeltz et. al., 2006). Rtn4a was initially characterized to insert specifically into peripheral ER tubules, and both Rtn4a and DP1 possess a hairpin domain that promotes membrane curvature, now called the Reticulon Homology Domain (RHD) (Voeltz et. al., 2006; Hu et al., 2008; Shemesh et al., 2014). Other proteins that include RHDs, but appear to have increasingly specialized roles in ER tubule network organization, are the small GTPase Atlastin and Lunapark (Chen et al., 2015; Wang et al., 2016). A separate class of proteins that possess RHDs are the Receptor expression enhancing proteins (Reeps). These family members were initially characterized to traffic certain GPCRs, such as olfactory receptors, to the plasma membrane but have since been demonstrated to function as key regulators of ER remodeling and microtubules (Voeltz and Prinz, 2007; Fan et al., 2022). Reeps are present in two major classes in most higher organisms: the Reep1-4 subfamily and the Reep5-6 subfamily. Less is known about what distinguishes the function of these individual Reeps outside of sequence and expression differences, and the Reep5-6 subfamily has been less characterized than the Reep1-4 subfamily (Björk et al., 2013).

Here, we use the syncytial *Drosophila* embryo as a model to investigate the roles of ER-shaping proteins during cleavage mitoses. Our findings indicate that *Drosophila* ReepB—a homolog of the Reep5-6 subfamily—plays a dominant role in organizing the ER during these divisions, as its disruption most significantly impairs mitotic progression. To further dissect how ReepB influences ER morphology, we employ an in vitro assay to examine its impact on ER dynamics. Our results suggest that different Reep subfamilies may vary in their shaping abilities on the ER, with ReepB likely contributing to the robust remodeling required for rapid processes such as the ER transitions observed during cleavage mitoses.

## Results

### A screen of ER shaping proteins reveals Receptor Expression Enhancing Proteins necessary for successful rapid mitoses

Previous work has shown that the ER and microtubule spindles maintain a close juxtaposition during mitoses in the *Drosophila* embryo, but it is unclear whether structural changes in ER morphologies modulate mitotic processes (Terasaki et al., 2003; Bobbinec et al., 2003; Bergman et al., 2015; Araújo et al., 2022; Rollins and Blankenship, 2023). Several families of ER tubule-associated proteins are present in the *Drosophila* genome. These include the Reticulon-like proteins (Rtnl1, Rtnl2), the Receptor Expression Enhancing proteins (ReepA, ReepB), Atlastin (Atl), and Lunapark (Lnpk).

As a starting point for this study, we screened each of these ER shaping proteins by shRNA-mediated disruption and scored division success during the rapid cortical divisions (syncytial cycles 10-13) (Figure 1A). UAS shRNA expression was driven by a maternal tubulin promoter in embryos co-expressing fluorescent Histone-2av (see Methods), and the subsequent cortical mitoses were monitored for division success (Figure 1A). In the first syncytial cycle (cycle 10), disruption of most ER shaping protein function (*Rtnl1, Rtnl2, atl, Lnpk,* and *ReepA*) had minimal impacts on mitosis (between 0-3% failed mitoses); however, *ReepB* disruption caused ∼20% of cycle 10 mitoses to fail completely (Figure 1A). These polyploid nuclei undergo fallout from cortically-associated regions and will no longer contribute to development (Figure 1B, D). Similarly, in the next cell cycle (cycle 11), *ReepB* disruption again caused three times the number of failed mitoses as the next closest ER shaping protein (Figure 1A). However, with each subsequent division, the rate of mitotic failures lessens in *ReepB* embryos – a trend that has also been observed in other backgrounds that disrupt ER function during the cleavage cycles (Rollins and Blankenship, 2023). By contrast, disruption of *Rtnl1, Lnpk,* and *ReepA* had an opposing trend, with failure rates increasing from cycle 10 to cycle 13 and peaking during cycle 13. Atlastin disruption produced milder defects than *ReepB* disruption but had a similar peak in the earlier cortical cycles (cycle 11) (Figure 1A), while *Rtnl2* disruption had little effect on divisions, consistent with high throughput transcriptional analysis that indicates that Rtnl2 is not significantly expressed at this stage of development (Brown et al., 2014).

**Figure 1:**
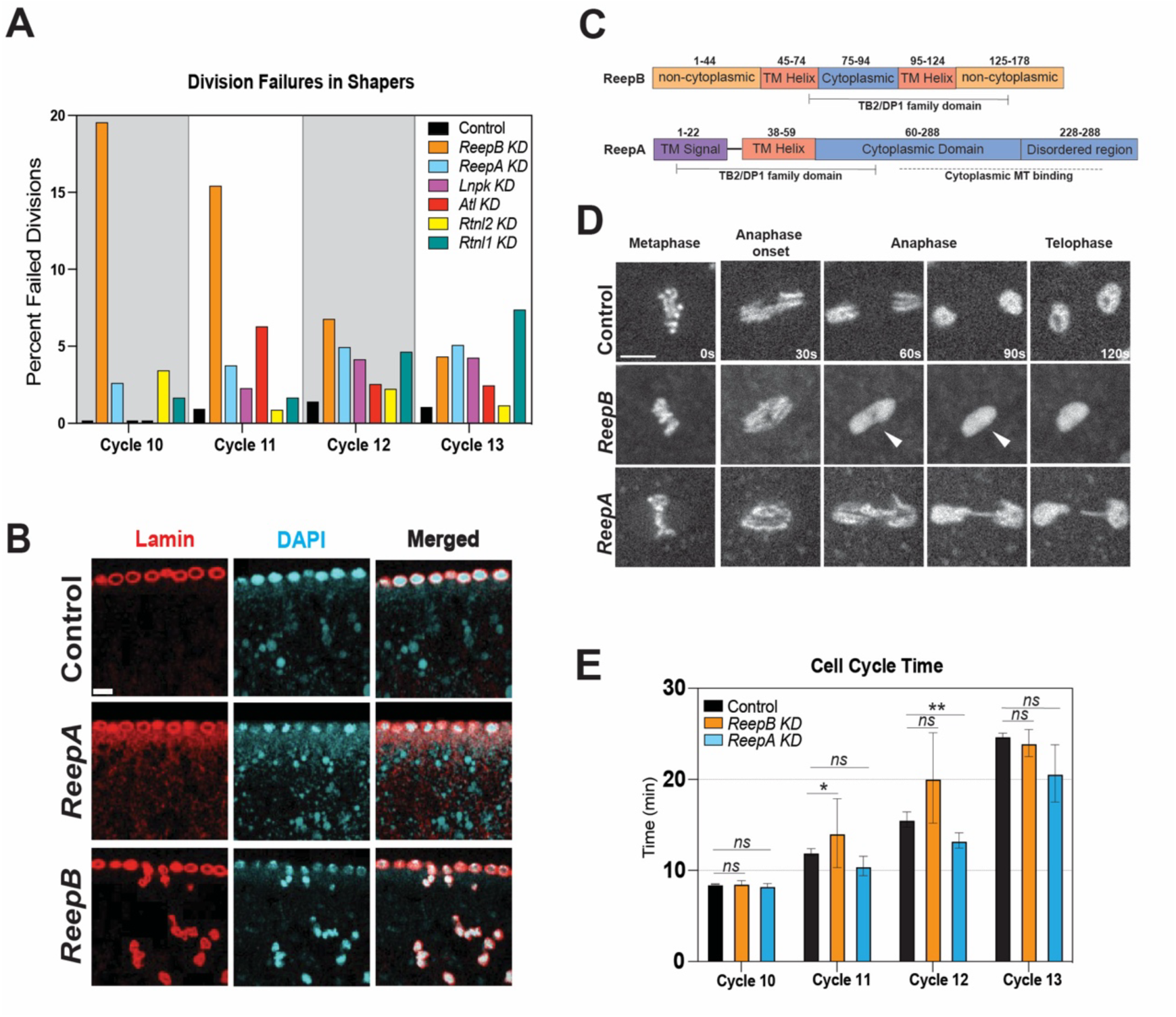
Screen of shRNAs targeting ER tubule shaping proteins in syncytial mitoses elucidates Reeps as primary mitotic ER shapers. **(A)** Quantification of ER shapers expressed in embryos with His2av marker to identify total percentage of division failure for each round of cortical cell divisions n≥90 divisions. **(B)** Nuclear fallout during division failure in ReepB knockdown. Scale bar = 5 μm. **(C)** Protein domain map of ReepA and ReepB. **(D)** Representative images of His2av in embryos expressing shRNAs targeted to *ReepB* or *ReepA* showing typical division failure phenotype at indicated mitotic phase. Arrowheads indicate collapsed mitosis. Scale bar = 5 μm. **(E)** Cell cycle time in each *Reep* depletion and control embryos for each round of cell division, n≥90 divisions. Statistics by Mann Whitney U-Test.

Given these screening results, we focused on the REEP family of proteins (ReepA and ReepB) to examine the effects of ER shaping and organization on cleavage mitoses. Examining ReepA and ReepB function may also provide data on how two shaping proteins from related sequence families are utilized during development (Figure 1C). Imaging mitoses after *ReepB* disruption revealed that the majority of these failures had defects characteristic of “spindle collapses” (Holly et. al., 2015), in which mitoses appear normal through metaphase and initiate anaphase but then fail as the separating chromosomes collapse back toward the metaphase plate (arrowheads in Figure 1D). By contrast, *ReepA* defective mitoses had a later cell cycle onset and lower prevalence (Figure 1A, D). These trends were further confirmed by creating independent, secondary shRNA lines which resulted in trends reflective of those above (Supplemental Figure 1). *ReepA and ReepB* disrupted embryos have similar cell cycle times as control embryos through the four syncytial blastoderm cycles, though *ReepB* cycle times trend longer in cycles 11 and 12, while *ReepA* can be slightly shorter than control in the last two syncytial cycles (Figure 1E). These data demonstrate that compromising the function of ER shaping proteins affects mitotic fidelity to varying degrees; however, certain proteins (ReepB and Atlastin), appear to have the strongest effects during cleavage-stage mitoses. Atlastin has been previously reported to effect spindle dimensions in the syncytial embryo (Araújo et al., 2022) and we therefore concentrated on Reep family function in our following analyses.

### ReepB depletion affects mitotic coat formation as excess sheeted ER surrounds the spindle

Given these results, we were interested in whether the ability to remodel ER morphologies around the inner mitotic space was compromised in *Reep* depleted embryos. To this end, we imaged embryos expressing a RFP:KDEL-ER marker (Figure 2A, B). Compromising *ReepB* function resulted in mitotic ER coated figures that form but possess massive deformations from typical ER morphologies as compared to control embryos (Figure 2A). Indeed, the ER envelope often appears to splay away from its normal juxta-spindle location creating angular bends in the ER coat (Figure 2A, arrowheads). Additionally, the mid-body accumulation of ER is absent in *ReepB* embryos. However, the arrangement of ER in *ReepA* disrupted embryos appears more like those formed in control embryos, while a fraction of mitotic figures still appear to have aberrant ER shapes (Figure 2A, B). Further examination of the ER in *ReepB* embryos at mid-mitotic stages shows ER “frills” that are highly characteristic of *ReepB* disrupted embryos and appear to radiate out from the coat (Figure 2B). Indeed, when analyzing the presence of ER abnormalities in either Reep depletion background, we observe that 92% of mitotic ER arrays display some level of disruption (deformation or frilling) in *ReepB* embryos, while 13% of *ReepA* ER morphologies at mitosis are notably perturbed (Figure 2C). Given the expected role of Reep proteins in promoting ER tubulation, it is likely that these phenotypes reflect a reduction in the ability to shape ER after Reep family disruption. This suggests that the aberrant ER morphologies after *ReepB* depletion may be due to an enrichment in ER sheets. Consistent with this, when imaging ER shapes in 3D, we find that these structures persist across multiple z-layers (Figure 2D). As the axial resolution of our confocal microscope is ∼500 nm and ER tubules are known to have a diameter of 30-60 nm, these structures are likely indicative of ER sheets forming where there would normally be greater tubule formation. These results suggest that ReepB is a key shaper of mitotic ER structures in the fast cortical divisions of the syncytium, and may also be a more potent ER shaper than ReepA at these stages.

**Figure 2:**
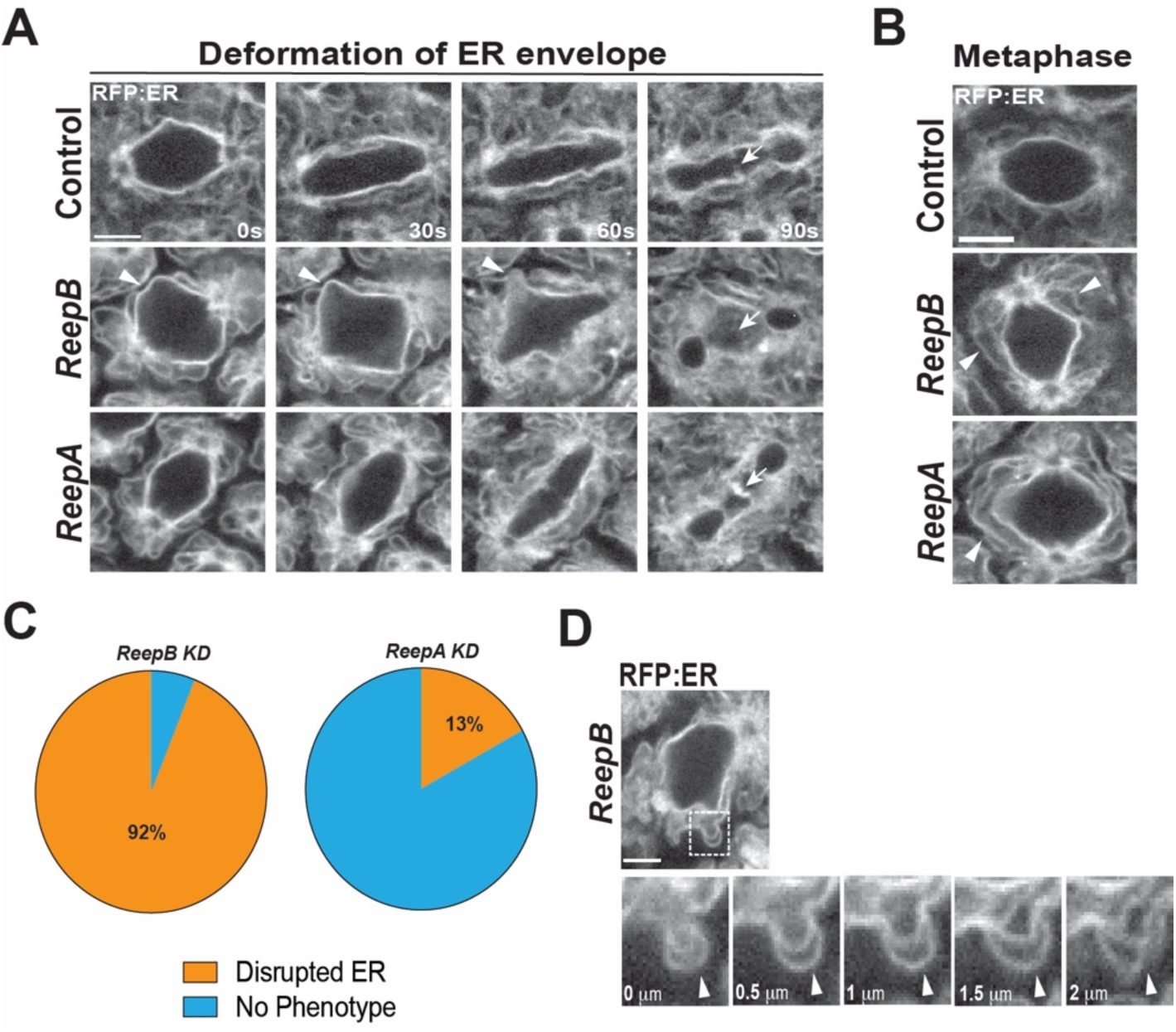
*ReepB* and *ReepA* depletion result in differentially penetrant ER shaping defects in which ER becomes more sheeted. **(A)** Still images from live imaging mitoses of embryos expressing UAS:RFP-KDEL and an shRNA targeted against *ReepB, ReepA,* or control (*white* gene). Arrowheads indicate perturbation of ER envelope shape, arrows mark midbody ER accumulation (or absence in *ReepB* compromised embryos). **(B)** Still images from live imaging of ER shapes during metaphase of cell cycle 11 in embryos expressing UAS:RFP-KDEL and shRNAs targeted against *ReepB* or *ReepA*. Arrowheads indicate formation of frilled, sheeted ER structures. Scale bars = 5 μm. **(C)** Qualitative analysis of ER disruption in *ReepB* and *ReepA* embryos in which either deformation of the ER envelope or disorganization of ER structures (A,B) is observed. **(D)** Z-scan of ER disorganized structures as previously shown in **B** in which continuity of signal is shown in Z direction indicating ‘sheet’ morphology. Scale bars = 5 μm.

### ReepB and ReepA are differentially expressed and non-redundant

One potential explanation for the greater impact of *ReepB* depletion at these stages is that ReepB may have been evolutionarily selected for use at these stages. Systematic high throughput analysis has indicated that ReepB is expressed at higher level at these stages, while ReepA expression strengthens once the embryonic epithelium has formed (Brown et al., 2014). To verify this, we performed quantitative PCR (qPCR) with primers for either ReepA or ReepB (Figure 3A). qPCR was performed in embryos collected within the syncytial stage (0-2 hrs) as well as embryos that were aged through this stage of development and into cellularization and gastrulation (2-4 hrs). Consistent with the modENCODE RNAseq database, we find that ReepB expression is elevated in early embryogenesis, and is then strongly reduced as further development occurs (Figure 3A, orange). By contrast, ReepA expression is reduced in earlier stages of embryogenesis (0-2 hrs) but increases in later stages (2-4 hrs; Figure 3A, blue). This results in ReepB expression being at significantly higher levels than ReepA in syncytial embryos, whereas the opposite is true for more developmentally advanced embryos. These differences are also apparent at the level of protein expression – semi-endogenous promoter constructs that have GFP tags incorporated at the C-terminus of either ReepB or ReepA also have much stronger levels of ReepB:GFP than ReepA:GFP during the cortical divisions (Figure 3B, C; see Methods for more construct information) (Yalçin et al., 2017). Taken together, these data suggest that ReepB is the developmentally dominant Reep and ER shaping protein at these stages, and may explain why *ReepB* disruption results in a higher degree of perturbation in syncytial embryos than when other ER shapers are compromised.

**Figure 3:**
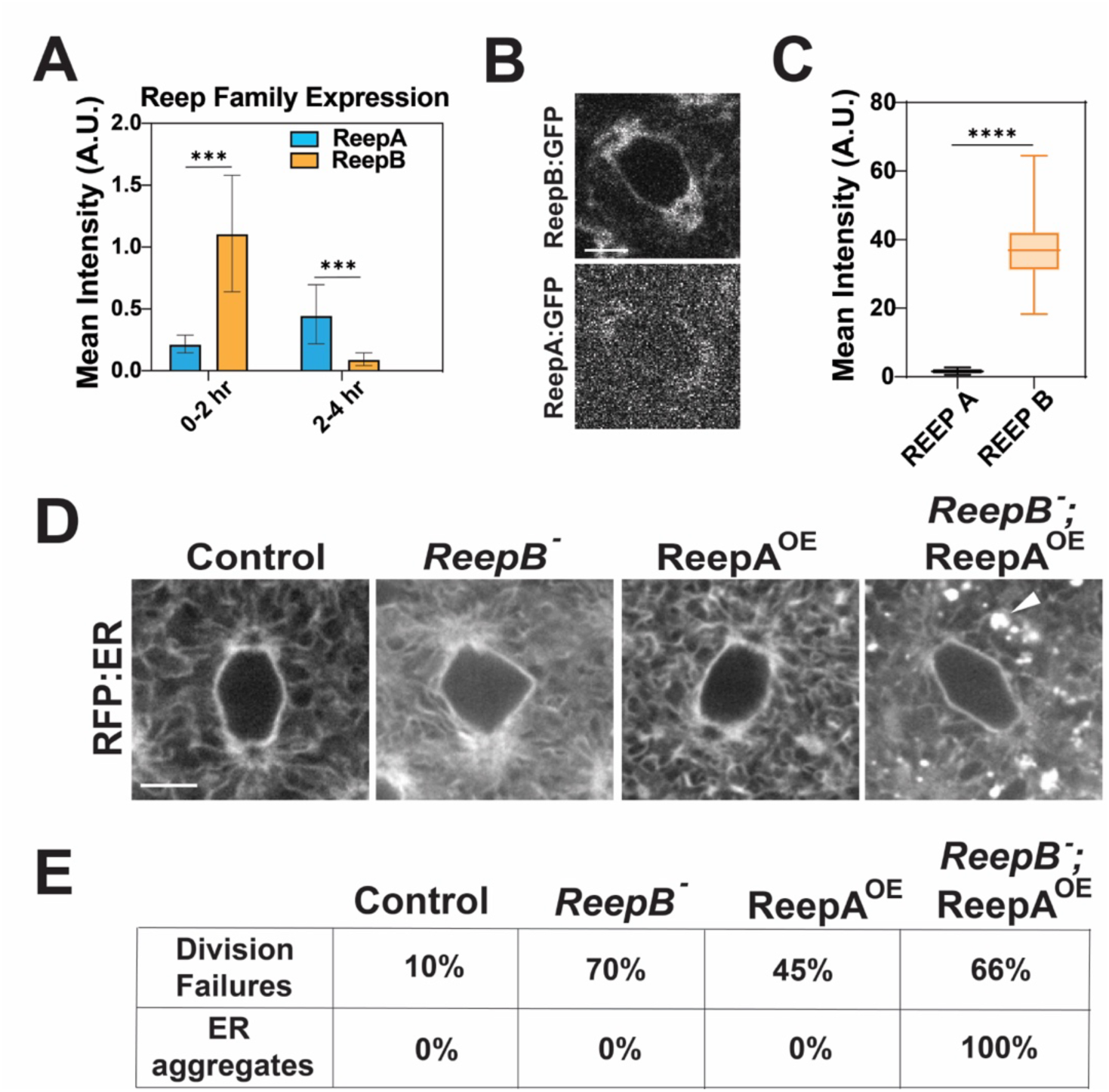
ReepB and ReepA are non-redundant and expressed in opposition. **(A)** Results from qPCR ΔΔCt analysis of ReepB and ReepA expression normalized to sqh in embryos at 0-2h post-oviposition (pre-syncytial and syncytial stage development) or 2-4h post-oviposition (cellularization through germband extension), n=4. **(B)** Still images from live imaging of ReepB:GFP and ReepA:GFP in cell cycle 11 at identical imaging parameters, n=40 measurements each. **(C)** Fluorescent intensity measurements of ReepB:GFP and ReepA:GFP in cell cycle 11. **(D)** Still images from live imaging of UAS:RFP-KDEL in control embryos, *ReepB* depleted embryos (*ReepB^−^)*, ReepA overexpressed embryos (ReepA^OE^), and *ReepB* depleted with ReepA overexpressed embryos in cell cycle 11. **(E)** Quantitation of phenotype penetrance in each condition from C based on the percentage of embryos displaying each phenotype. Scale bars = 5 μm. Statistics by Mann Whitney U-Test.

To further explore this dynamic, we attempted to rescue *ReepB* depletion by overexpressing ReepA in syncytial embryos. ER (KDEL:ER marker) dynamics were imaged in embryos that overexpressed ReepA along with *ReepB* shRNAs (Figure 3D). We also examined embryos overexpressing ReepA without *ReepB* disruption to determine whether ReepA overexpression alone can perturb ER structures. Interestingly, ReepA overexpression embryos have a high level of division failures (∼45% at cycle 11), suggesting that the relative lack of expression of ReepA at these stages is a critical element in the cortical divisions (Figure 3D, E). However, little observable impact on ER morphologies was apparent in this background at the level of in vivo fluorescent imaging (Figure 3D, E). When ReepA is overexpressed alongside *ReepB* disruption, division failures are enhanced, and, intriguingly, ER structures appear to be highly perturbed (Figure 3D, E). ER is often observed to accumulate in dense aggregates and the spindle-associated ER is weakened (Figure 3D arrowheads, and E). These data indicate that ReepA and ReepB are not simply redundant in their function but suggest that specific activities are associated with each protein that are appropriate for specific developmental times and processes. It is also interesting that a new ER aggregation phenotype occurs when the “wrong” Reep (ReepA) is expressed in the absence of the syncytial Reep (ReepB), suggesting that ReepB may possess a protective activity to prevent the formation of these aggregates. Given these effects, we wanted to identify how ReepB acts in the cortical divisions to allow the fast mitoses that are prevalent at these stages.

### ReepB disruption affects spindle size and dynamics during early division cycles

Given the above defects in mitoses, we examined whether spindle function is affected by the aberrant ER envelope and sheet structures observed after Reep disruption. To this end, we imaged spindle dynamics by endogenously labeling microtubules with the MT-associated protein, Jupiter:GFP. Although gross spindle morphologies are relatively normal after *ReepA* or *ReepB* depletion, these data revealed that at metaphase stages spindle dimensions are reduced in *ReepB* disrupted embryos (Figure 4A-C). Cycle 11 *ReepB* spindle widths are reduced by 1.8 μm on average, while spindle lengths decrease by ∼2 μm. *ReepA* disruption does not appear to affect spindle sizes in either dimension, consistent with the low level of mitotic defects in this background (Figure 4B, C). We also analyzed whether spindle elongation, which helps drive anaphase behaviors, was similarly affected. Indeed, late metaphase and anaphase spindle elongation is defective in *ReepB* shRNA embryos, and these defects are not present in *ReepA* shRNA embryos (Figure 4D-G). Interestingly, the onset of ER phenotypes (including disorganized sheet structures and deformed spindle envelopes) were observed as early as prometaphase and metaphase respectively (Figure 3D), potentially consistent with a causative relationship between defects in spindle elongation and the effects that ReepB disruption have on ER organization and shaping. These phenotypic defects in spindle structure may underlie the division failures that are seen in *ReepB* embryos.

**Figure 4:**
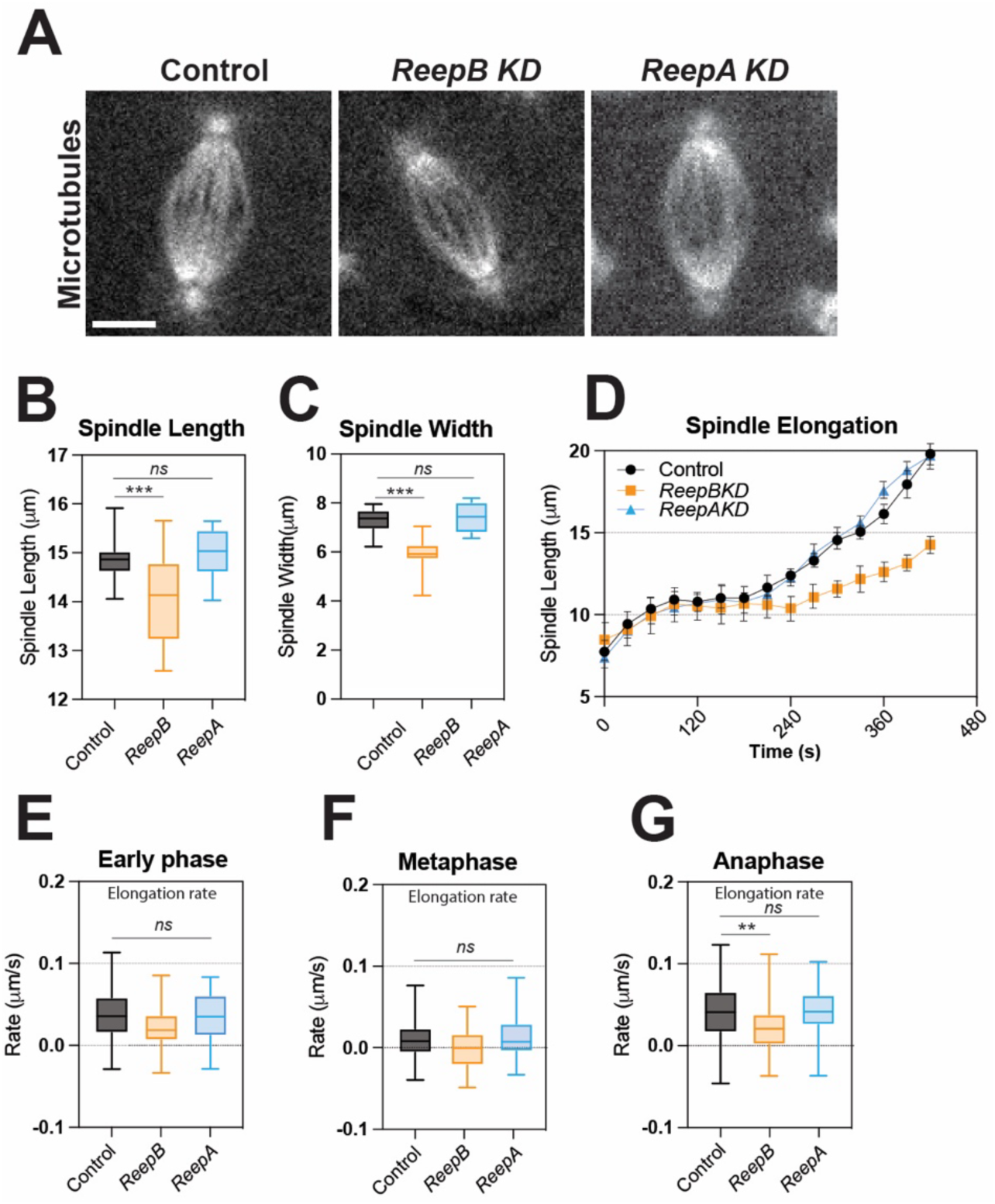
*ReepB* depletion affects spindle size and elongation during mitotic metaphase and anaphase. **(A)** Still images from live imaging of Jupiter:GFP protein trap, a microtubule binding protein, in control embryos or embryos expressing shRNAs target to ReepB or ReepA. Scale bar = 5 μm. **(B)** Measurement of spindle length in cell cycle 11 during metaphase in each condition from A, n≥112. **(C)** Measurement of spindle width, measured across the metaphase plate in each condition, n≥112. **(D)** Measurement of spindle length overtime in each condition at 20s intervals in cell cycle 11 displaying a lag of spindle elongation after ∼240s. n=15. **(E)** Spindle elongation rates in early cell cycle (0-100s from centrosome opposition). **(F)** Spindle elongation rates during metaphase pre-active elongation in cell cycle 11 (100-220s). **(G)** Spindle elongation rates in anaphase in each condition during cell cycle 11 (220-440s). Statistics by Mann Whitney U-Test.

A marked reduction of microtubule densities within the syncytial spindles is correlated with failed divisions (Rollins and Blankenship, 2023). To further determine whether the anaphase spindle is robust enough to separate the daughter chromosomal complements, we analyzed the spindles based upon microtubule intensity (Figure 5A-C). We find that microtubule (MT) spindle intensities are decreased relative to controls as early as metaphase. Notably, this correlates temporally with the defects in spindle length and elongation rates in *ReepB* embryos. MT intensities in *ReepB* embryos begin to deviate from controls at ∼150-200 s into the cell cycle, whereas elongation rate differences were also seen by 200 s (Figures 4D, 5B). By anaphase, *ReepB* embryos display a ∼27% loss of MT intensities compared to control embryos (Figure 5C) suggesting the perturbed ER spindle envelope may affect microtubule densities. We were also interested if these defects could be observed at the level of critical MT nucleators such as γ-tubulin. Intriguingly, in *ReepB* and *ReepA* depleted embryos we find that γ-tubulin is reduced compared to controls at mitotic time points (41% decrease in *ReepB* and 21% decrease in *ReepA* embryos from controls), while interphase γ-tubulin is near control levels (Figure 5D-F). These results are suggestive of a possible link between ER structure and the remodeling of spindle and MT function necessary for the separation activities that occur at anaphase. Indeed, MT spindles fail to maintain contact with the ER envelope in *ReepB* compromised embryos (Figure 5G). Given these data, we next wanted to examine whether the two *Drosophila* Reeps function differently in regulating ER tubular morphologies and how these differences may impact spindle scaling.

**Figure 5:**
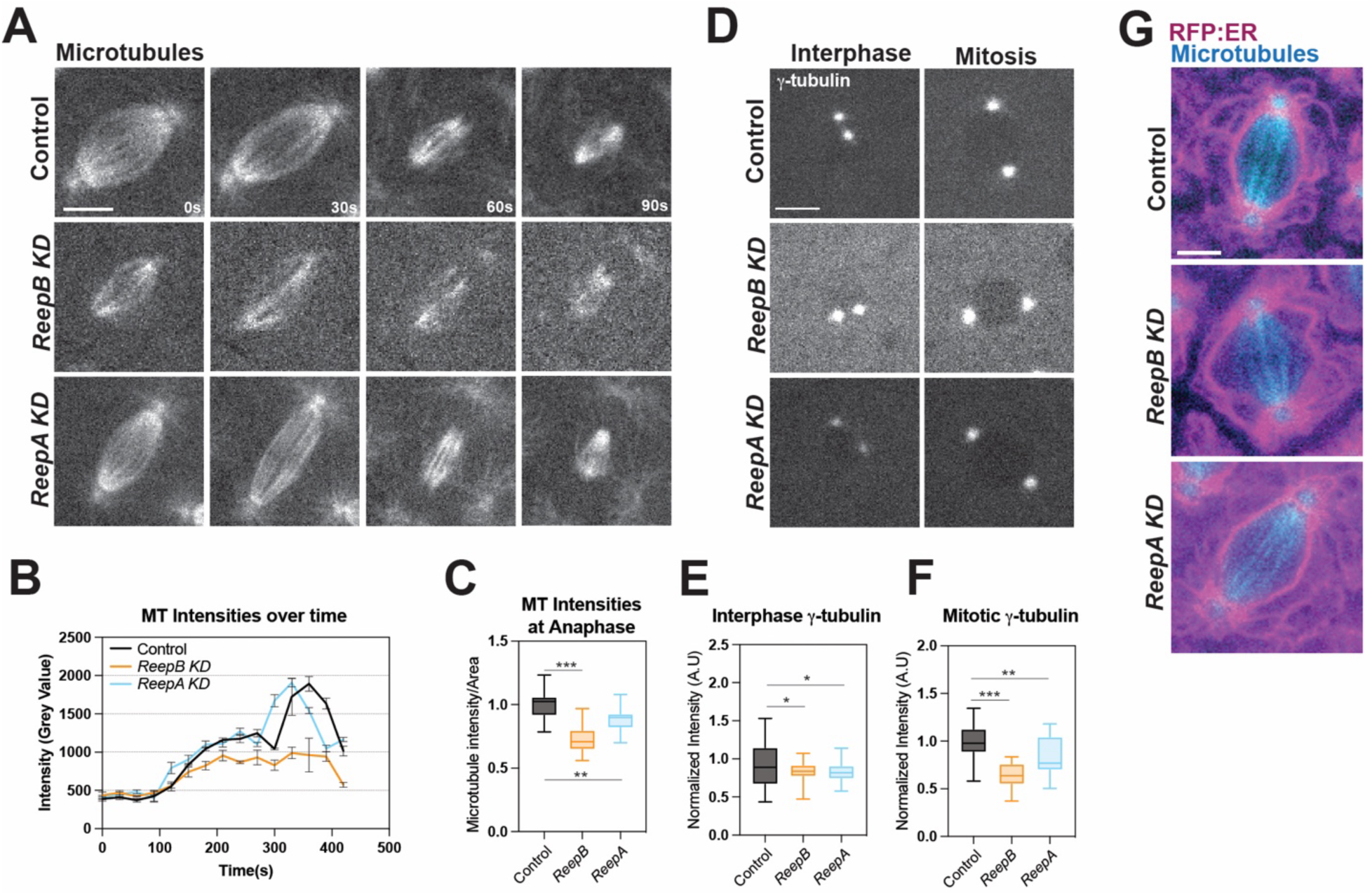
Anaphase spindles display reduced microtubule intensities and γ-tubulin presence. **(A)** Still images from live imaging of embryos expressing shRNAs targeted to either *ReepB* or *ReepA* as well as non-shRNA expressing sibling controls in conjunction with Jupiter:GFP protein trap (Microtubules) at 30s intervals in cell cycle 11 beginning after the formation of a metaphase spindle. **(B)** Quantification of microtubule intensities in the spindle measured from identically leveled and photobleach-corrected movies at cycle 11. n≥15 spindles per condition. **(C)** Microtubule intensities measured at late anaphase to display endpoint differences in MT intensities. n≥81 spindles per condition. **(D)** Still images from live imaging of γ-tubulin:GFP in control embryos or those expressing shRNAs targeted to either Reep in interphase (after centrosome duplication) and mitosis (prometaphase). **(E)** Quantification of γ-tubulin in interphase of cell cycle 11 of all 3 conditions, n≥130 measurements. **(F)** Quantification of γ-tubulin in mitosis of cell cycle 11 of all 3 conditions, n≥130 measurements. **(G)** Still images from live imaging of RFP:KDEL-ER and Jupiter:GFP protein trap in cell cycle 11 in controls or embryos expressing shRNAs target to either Reep. Scale bars = 5 μm. Statistics by Mann Whitney U-Test.

### Expression of Drosophila Reeps in Cos-7 cells to examine tubular networks

Our results to this point illustrated *in vivo* differences in the requirements and function of the two *Drosophila* Reeps – ReepA and ReepB. However, it is difficult to resolve tubular ER morphologies in the early embryo by either live or fixed imaging. To visualize ER networks with high resolution, we engineered expression vectors for each *Drosophila* Reep tagged with a C-terminal GFP and transfected them into Cos-7 cells along with empty vector controls. Cells were co-transfected with Halo-Sec61β as an ER marker and visualized at 15 hours post-transfection. We observed that expressing *Drosophila* Reep proteins did not grossly affect ER tubule networks and that both Reep constructs localized to ER tubules, while the control vector showed a cytoplasmic GFP signal (Figure 6A). ReepB-GFP localized strongly to tubular ER (Pearson R=0.972 ± 0.22), while ReepA-GFP colocalization with tubules was weaker (Pearson R=0.526 ± 0.19) (Figure 6B-D). Given these different relative levels of association with the tubular ER, we wanted to analyze whether the movement of ER tubules was differentially affected when comparing these two conditions. Indeed, when live-imaging cells over short time frames, we find that cells expressing ReepB-GFP display greater rates of tubule elongation compared to control cells or ReepA-GFP expressing cells (Figure 6E, F). When ReepA-GFP is overexpressed, we also saw an increased rate of tubular elongation compared to controls. However, this is a more moderate increase than was observed after ReepB-GFP expression (Figure 6E, F). Finally, retraction rates across all three conditions were not significantly different in our observations, suggesting Reeps have a stronger influence on the formation and elongation of ER tubules, but not their shortening (Figure 6G). Given these results, we were curious if we could detect accumulations or movements of the ReepB or ReepA tagged proteins within cells at elongating tubule tips. To this end we used our live imaging data to analyze Reep enrichment relative to Halo-Sec61 enrichment. Interestingly, we find that Halo-Sec61 increases at an elongating tip later than ReepB-GFP, although this trend is not seen in ReepA-GFP cells (Figure 7A, B). Taken together, these results show that the expression of Reep family proteins can have differential impacts on the elongation of ER tubules – ReepB expression appears to more potently affect tubule elongation than ReepA. These results are consistent with our data in the *Drosophila* embryo, as depleting ReepB resulted in an ER that was less able to adapt to the changes required for syncytial mitoses, suggesting that ReepB may be a potent ER shaper specifically recruited for rapidly changing ER morphologies.

**Figure 6:**
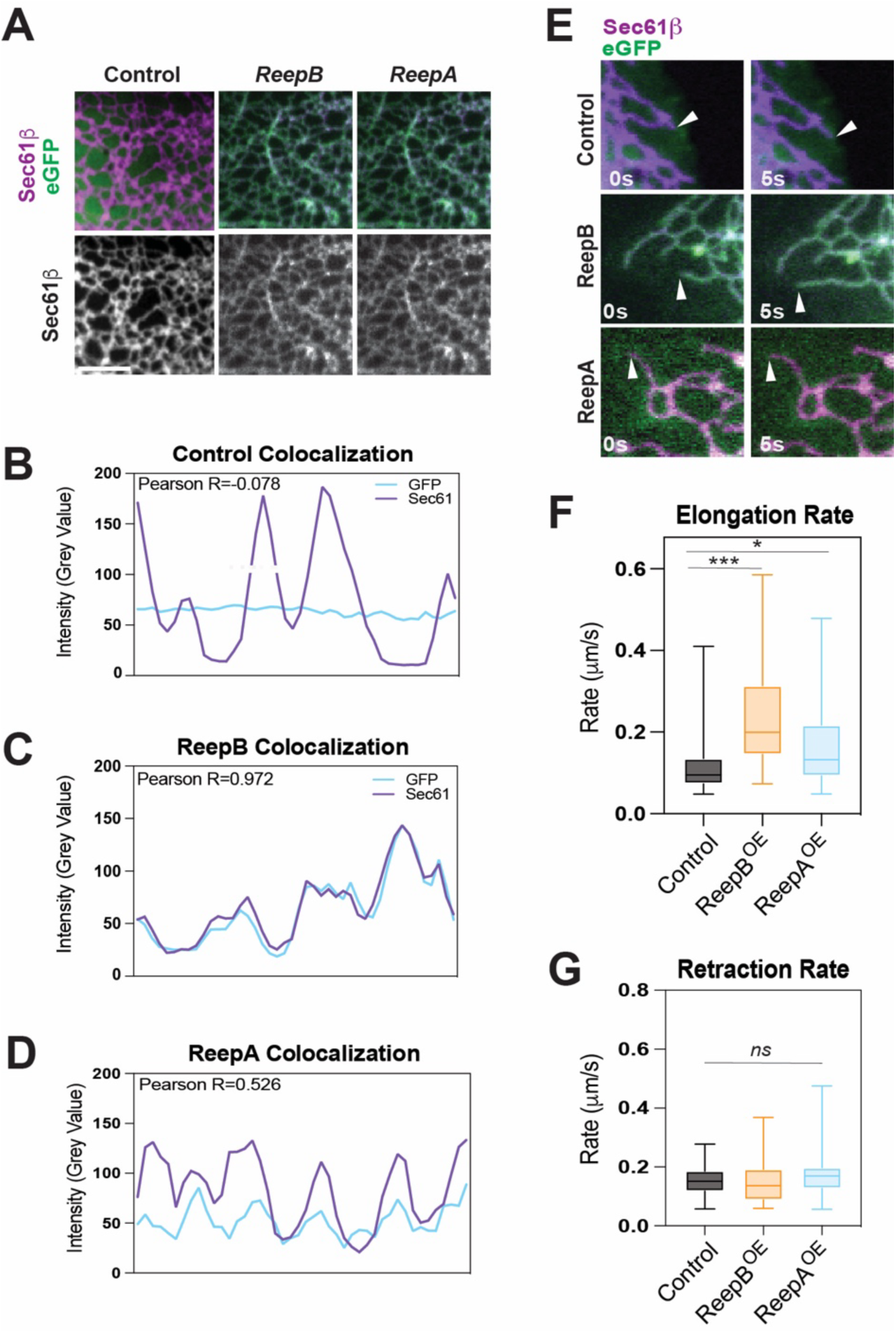
Expression of ReepA and ReepB in Cos7 cells to examine tubular dynamics. **(A)** Still images from live imaging of Cos7 cells transfected with Halo-Sec61β and pEGFP-N1 vector with ReepB, ReepA, or empty vector for control. Scale bars = 5 μm. **(B-D)** Representative line scans of ER tubule and GFP intensities in each condition in control cells (Halo-Sec61 and vector control, ReepA-pEGFP, or ReepB-pEGFP. Statistics by Pearson analysis. n=15. **(E)** Still images from live imaging of Halo-Sec61β and ReepA-GFP or ReepB-GFP expressed in Cos7 cells at two time points. Arrowheads indicate the terminal end of an elongating ER tubule. **(F)** Quantification of the elongation rate of ER tubules in Cos7 cells in control, ReepA-GFP and ReepB-GFP cells. n≥52 measurements. **(G)** Quantification of the retraction rate of ER tubules in Cos7 cells of each condition. n≥46 measurements. Scale bar = 6 μm. Statistics by Mann Whitney U-Test.

**Figure 7:**
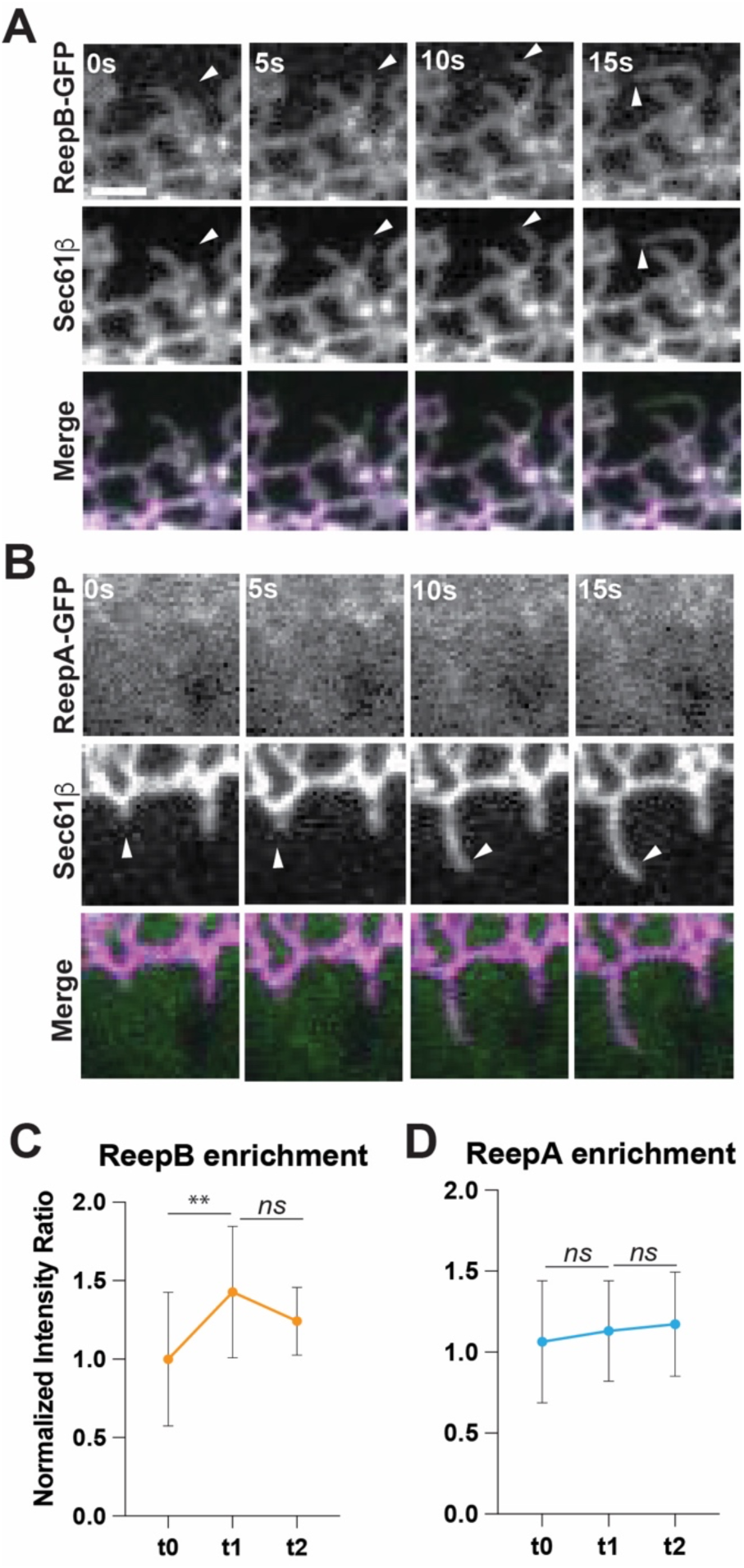
ER tubules undergoing elongation in Cos7 cells show an enrichment of ReepB relative to Sec61. **(A,B)** Still images from live imaging of Cos7 cells overexpressing ReepB-GFP (A) or ReepA-GFP (B) and Halo-Sec61β at 5s time intervals in which a tubule is actively elongating. Arrowheads indicate terminal tip of an elongating ER tubule. **(C)** Quantitation of ReepB-GFP/Sec61β at the terminal tip of elongating ER tubules normalized to the initial time point. Pre-elongation marked by “t0”, active elongation marked by “t1”, stabilized tubule formation marked by “t2”. n≥25 measurements. **(D)** Quantitation of ReepA-GFP/Sec61β at the terminal tip of elongating ER tubules normalized to the initial time point. Pre-elongation marked by “t0”, active elongation marked by “t1”, stabilized tubule formation marked by “t2”. n≥22 measurements. Scale bar = 2 μm. Statistics by Mann Whitney U-Test.

## Discussion

In this work, we examined the impact of ER morphology on microtubules and spindle function. The observed defects in the mitotic separation machinery after ER shaping disruption, in turn, undermined genomic segregation and stability. This is also consistent with the dramatic remodeling of ER structures that occur during mitosis. Indeed, screening various ER tubule promoting proteins, including Rtnl1, Rtnl2, Atlastin, Lunapark, ReepA and ReepB, revealed a main tubule shaper that is required during the rapid cleavage mitoses in the cortical syncytium. Disrupting *ReepB* through shRNA targeted knockdown, resulted in increased division failures that were present in earlier mitotic cycles (cycle 10 and 11), but self-rescued in later cell cycles. Interestingly, this self-rescue trend in division failure is reflective of those seen when the ER over-accumulates in embryos with Rab1 GTPase disruption (Rollins and Blankenship, 2023). One possible explanation for this early sensitivity to ER malfunction may be that cell cycles 10 and 11 (∼9-11 mins duration for cycles 10 and 11) are more rapid than cycles 12 and 13 (∼14-18 mins for cycles 12 and 13) (Xie and Blankenship, 2018). These later cycles may have additional time for ER remodeling to occur, thus allowing for other shapers (or residual function) to adapt the malformed ER coat of *ReepB* depleted embryos. Alternatively, the amount of ER (as measured by KDEL reporter intensities) associated with each nucleus is depleted after each round of division (Rollins and Blankenship, 2023). Thus, the absence of proper shaping may be less deleterious in the later cycles with smaller amounts of ER.

Interestingly, the malformation of ER envelopes in *ReepB* depleted embryos is associated with division failures and mitotic figures with reduced spindle dimensions in both length and width. Additionally, microtubule intensities also decreased in the spindle. Lastly, the ER coat around the spindle often shows gaps and bulges away from the inner spindle space. This suggests a potential link between ER contact sites and the spindle machinery which is necessary for the proper assembly of spindle microtubules during mitosis. Importantly, these phenotypes are *ReepB* specific. *ReepA* depleted embryos do not display, or only weakly possess, similar defects despite being a closely related protein. Indeed, overexpressing ReepA in *ReepB* depleted embryos generated even deeper disruptions in ER morphologies, with large, dense aggregates forming. Thus, ReepA does not appear to be able to substitute for ReepB function at these stages, consistent with RNA and protein expression profiles in which ReepB is much more strongly expressed in the early embryo. Studies from mammalian cells have suggested that the ER transforms to higher curvature and more tubulated structures during mitosis, and Reep family proteins have been implicated in this transformation (Pukha et al., 2007; Pukha et al., 2012; Schlaitz et al., 2013; Kumar et al., 2019). Our *in vivo* results support these findings, and further implicate this transition in feeding back on the mitotic spindle maturation and function.

How might the endoplasmic reticulum affect spindle formation? ER proteins such as STIM1 and TAOK2 have been implicated in mediating ER interactions and binding microtubules (Grigoriev et al., 2008; Nourbakhsh et al., 2021), but disruption of these proteins in the early embryo did not affect the cleavage divisions (our unpublished data). Alternatively, REEP4 has been suggested to directly bind microtubules (Schlaitz et al., 2013). This function was linked to a pathway that mediated the clearance of ER from metaphase chromatin, but it may be that Reep family proteins use this microtubule binding capacity to stabilize and/or organize spindle microtubules. Other possible requirements for Reep shaping function could involve the necessity for the ER to absorb and shape the large surface area of the nuclear envelope after mitotic breakdown, or that tubulated regions of the ER are more active in lipid synthesis (reviewed in Schwarz and Blower, 2015). The direct mechanism for how defects in ER shaping impact spindle morphologies is not clear, but further work will be needed to define why ER tubulation (or the high-curvature domains that Reep proteins generate) are necessary for robust spindle assembly.

## Methods

### Fly Stocks and Genetics

Fly stocks used in this study are as follows: UAS-RFP:KDEL III (BL-30910), His2Av:RFP III (BL-23650), Jupiter:GFP Protein Trap (BL-60156), and ncd-γ-tubulin:GFP (BL-57328), ReepA Val20 (BL-37500), ReepB Val20(BL-62476), Lunapark Val20 (50998), Atlastin Val20 (BL-36736), Rtnl2 Val20 (BL-58208), pUAS:ReepA:FLAG (BL-93604), M{ReepB-GFP}ZH-86Fb (BL-77907), M{ReepA-GFP}ZH-86Fb (BL-77906). M{ReepB-GFP}ZH-86Fb (BL-77907) and M{ReepA-GFP}ZH-86Fb (BL-77906) are P[acman] clones of CH322-97D15 derived from BAC containing genomic regions around ReepA/B and recombineered into attP2 on chromosome 3 and thus should be expressed by near-endogenous promoter elements (Yalçın et al., 2017). BL stocks were all obtained from Bloomington Drosophila Stock Center. Additionally, Resille:GFP (A. Spradlin, Carnegie Institution), ReepB Walium22 primers, ReepA Walium22 primers, and Rtnl1 Walium22 primers were all generated using the Drosophila Research and Screening Center protocol (https://fgr.hms.harvard.edu/knockdown-vectors). Primers were annealed and cloned into pWalium22 vector (DGRC Stock 1473; https://dgrc.bio.indiana.edu//stock/1473 ; RRID:DGRC_1473) to create RNAi lines targeted to each gene of interest. The construct was then verified through DNA sequencing. All fly crosses were maintained at 25°C. Rtnl1wal22 expresses an shRNA targeting the sequence CTACGAGAACAACAAGCAA. ReepBwal22 expresses an shRNA targeting the sequence GTTCTACCATCATCTACAA. ReepAwal22 expresses an shRNA targeting the sequence GACCAAGGATGTCAAGGAA. Phenotypes were confirmed in both Walium and Valium lines. UAS flies were crossed to matαTub-Gal4VP16 67C;15 (D. St. Johnson, Gurdon Institute, Cambridge, UK) females to drive transgene expression.

### Live Imaging

Live imaging of embryos was carried out on a CSU10b Yokogawa spinning-disk confocal microscope from Zeiss and Solamere Technologies. Embryos were collected on apple juice agar plates with yeast and dechorionated in 50% bleach solution, washed with deionized water, then transferred to a slide with a gas-permeable membrane in Halocarbon 27 oil (Sigma-Aldrich). A cover slip was placed over the embryos for imaging. All images were obtained with a 63×/1.4 NA objective except for images of microtubules obtained for measuring spindle lengths which were obtained with a 100×/1.25 NA objective. Images for initial characterization of phenotypes were obtained in 25-layer z-stacks at a 30 second imaging interval. Images used for precise time-resolved measurements were obtained in 2-5 slice z-stacks at 1-5 second intervals. All z-stacks were taken at 0.5µm z-intervals between slices.

### Embryo fixation and immunostaining

Embryos were dechorionated with 50% bleach then washed with deionized water and manually devitellinized. Collected embryos were fixed at the interface of n-heptane and 4% formaldehyde in 0.1M sodium phosphate buffer (pH 7.4) for 1 hour. Embryos were then manually devitellinized and incubated with mouse anti-Lamin (DSHB #ADL, 1:100) and DAPI. Primary antibody stains were carried out overnight in 4°C with gentle agitation. Secondary antibodies conjugated with Alexa 488 or Alexa 568 (Molecular Probes, #A11034, #A11031) were used at 1:400 dilution for 45 minutes. Secondary stains were carried out at room temperature. Embryos were mounted on slides with Prolong Gold Antifade reagent +DAPI (Invitrogen, P36935) before imaging. Images of immunostained embryos were captured on an Olympus Fluoview FV1000 confocal laser scanning microscope with a 60×1.35NA objective.

### Cloning of Reep Overexpression vectors

Semi quantitative PCR analysis of *Drosophila* ReepA and ReepB isoforms was first performed to select for the dominant isoform in syncytial stages of embryogenesis by extracting RNA from embryos. Reverse transcription of RNA samples was performed using the Qiagen QuantiTect Reverse Transcription Kit (Qiagen #205311). Dominant isoform sequences were then cloned into the pEGFP-N1 vector (Addgene #172281). ReepB was inserted into the PstI-KpnI insertion site and ReepA was inserted into the EcoRI-KpnI site. A 12 amino acid linker (GSSG_n_) was inserted between the Reep sequence and the eGFP of the plasmid and DNA was amplified and isolated. Sanger sequencing was utilized to verify plasmids used in transfection.

### COS-7 cell culture and transfection

COS-7 (ATCC #CRL-1651) were maintained at 37°C and 5% CO_2_ in complete growth medium consisting of 10% fetal bovine serum (Gibco #A52567-01), 2 mM L-glutamine (Gibco #25030-081), and 1x Penicillin-Streptomycin (Gibco #15140-122) in DMEM (Gibco #11054-020). 25 mm #1.5 glass coverslips (Warner Instruments #CS-25R15) were cleaned by heated sonication in 1.5-2M potassium hydroxide solution followed by Hellmanex III (Sigma-Aldrich #Z805939), ddH_2_O, and stored in 95% ethanol. Prior to use, coverslips were coated with human fibronectin (Millipore Sigma #FC010) in 1x DPBS (Gibco #14200-075) for 30 minutes. COS-7 cells were plated to fibronectin coated coverslips and allowed to settle prior to PEI-based transfection of a total of 2 μg of DNA. Cells were transfected with 1.5 μg of either pEGFP-N1, pReepA-EGFP-N1, or pReepB-EGFP-N1 and 0.5 μg of Halo-Sec61-C18 (Addgene #123285).

### COS-7 cell imaging

COS-7 cells were imaged 12h, 15h, or 24h post transfection. Prior to imaging, cells were costained with 5 μM DRAQ5 (ThermoFisher #62251) and 1.25 μM HaloTag Ligand TMR (Promega #G8252) for 10 minutes followed by a 30-minute wash. All samples were imaged at 37°C and 5% CO_2_ in FluoroBrite DMEM (Gibco #A18967010). Imaging was done utilizing a Hamamatsu EM-CCD camera and a Nikon Eclipse Ti inverted microscope with a Yokogawa CSU-X1 spinning disk system using a 63X NA 1.40 Nikon oil objective lens (system built by Solamere Technologies). Cells were imaged at 1-minute intervals for 16-30 minutes with monitoring to check for phototoxicity. For fast resolution of peripheral ER growth and retraction, cells were imaged at 5s intervals for 5-8 minutes total and monitored to check for phototoxicity.

### Intensity measurements and normalization

Intensity measurements were obtained from live imaging with spinning disk confocal microscopy or from fixed and stained embryos (laser-scanning confocal). Embryos for intensity comparisons were imaged under identical imaging conditions to their controls (laser power, gain settings, exposure, etc.). In each imaged cycle, comparisons were set to the same z-layer, developmental time point, and fluorescence leveling as its respective control before obtaining intensity measurements. For intensity measurements, a region of interest was traced with freehand line or freehand area tool on ImageJ. ImageJ measurements of area, mean intensity values, max intensity values, and min intensity values were taken for each trace. Background fluorescence was analyzed by tracing and obtaining measurements for a similar cytoplasmic length/area within the same z-layer and at the same time point. Five background measurements were averaged together, and average background was subtracted from the measured intensity. Obtained intensity values were then divided by mean intensity of controls to obtain a normalized value of fluorescence.

### Area and length measurements

Images for obtaining area measurements (compartmental and ER size, for example) were measured with the freehand selection tool on ImageJ. The perimeter of structures was traced, and areas measured in pixels. Pixel areas were converted to micron areas based on the pixel resolution of the microscope and objective used. For spindle lengths, the line tool was used to draw a single line across the spindle from pole to pole based on apparent centrosome location.

### Statistics and Repeatability

All measurements represented in figures were obtained from imaging data of at least three embryos of at least three individual trials. In figure captions, k represents the number of embryos and n represents the number of individual measurements. A Mann-Whitney U-Test was used as denoted in figure legends. ns= not significant; * = p<0.05; **= p<0.005; ***= p<0.0005. Error bars indicate standard error.

### Image Editing and Figure Preparation

Images obtained through spinning disk and laser scanning microscopy were edited using Adobe Photoshop. Graphs were developed with Prism and Figures were created in Adobe Illustrator.

## Acknowledgements

We thank members of the Blankenship Lab for reading and providing constructive commentary of the manuscript. This work was supported by grants from the National Institutes of Health and NIGMS R01GM141243 and to JTB. Stocks obtained from the Bloomington Drosophila Stock Center (NIH P40OD018537) were used in this study and are gratefully acknowledged.

---

**Supplemental Figure 1:**
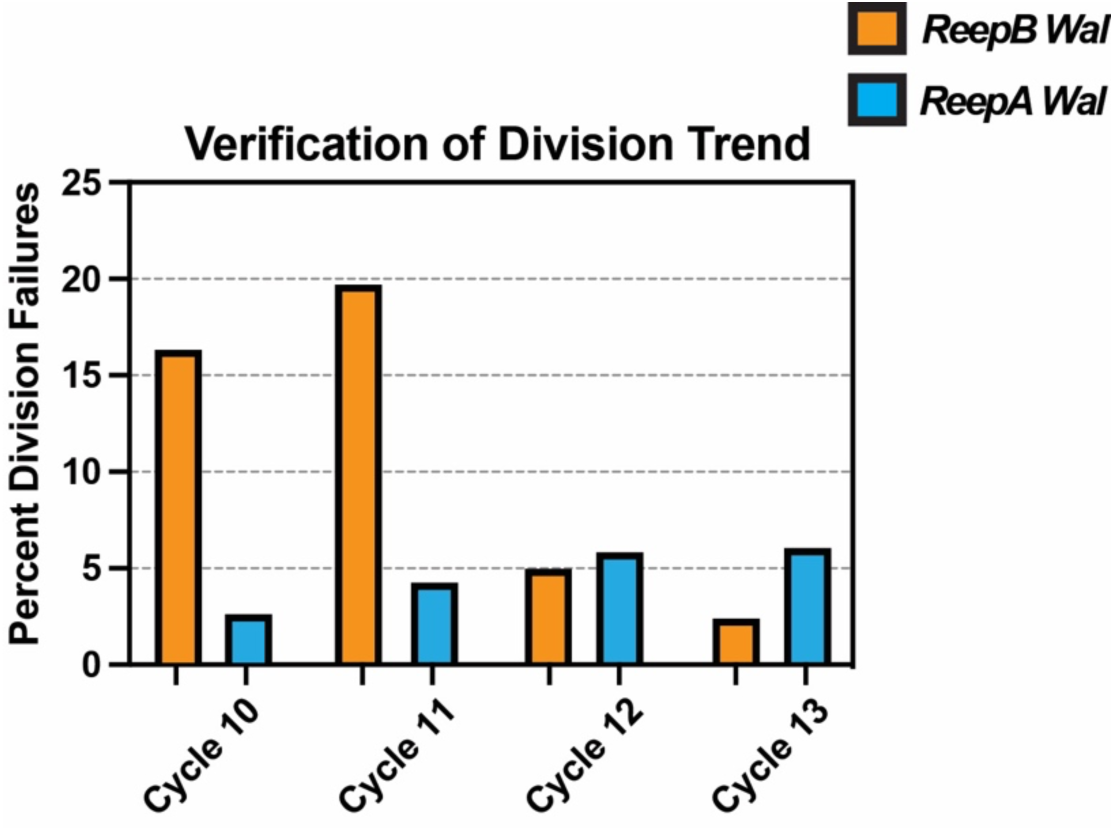
Independent shRNA lines demonstrate similar defects in mitoses during syncytial cycle. shRNA lines against ReepA and ReepB created in pWalium22 with unique targeting sequences from constructs reported in Figure 1.

